# Meta-Reinforcement Learning reconciles surprise, value, and control in the anterior cingulate cortex

**DOI:** 10.1101/2024.05.15.592711

**Authors:** Tim Vriens, Eliana Vassena, Giovanni Pezzulo, Gianluca Baldassarre, Massimo Silvetti

## Abstract

The role of the dorsal anterior cingulate cortex (dACC) in cognition is a frequently studied yet highly debated topic in neuroscience. Most authors agree that the dACC is involved in either cognitive control (e.g. voluntary inhibition of automatic responses) or monitoring (e.g. comparing expectations with outcomes, detecting errors, tracking surprise). A consensus on which theoretical perspective best explains dACC contribution to behaviour is still lacking, as two distinct sets of studies report dACC activation in tasks requiring surprise tracking for performance monitoring and cognitive control without involving surprise monitoring, respectively. This creates a theoretical impasse, as no single current account can reconcile these findings. Here we propose a novel hypothesis on dACC function that integrates both the monitoring and the cognitive control perspectives in a unifying, meta-Reinforcement Learning framework, in which cognitive control is optimized by meta-learning based on tracking Bayesian surprise. We tested the quantitative predictions from our theory in two different functional neuroimaging studies at the basis of the current theory crisis. We show that the meta-Reinforcement Learning perspective successfully captures all the neuroimaging results by predicting both cognitive control and monitoring functions, resolving the theoretical impasse about dACC function within an integrative framework. In sum, our results suggest that dACC function can be framed as a meta-learning optimisation of cognitive control, providing an integrative perspective on its roles in cognitive control, surprise tracking, and performance monitoring.

**Significance statement:** An important debate in cognitive neuroscience concerns the role of the anterior cingulate cortex (ACC) in cognition. Two effective and competing frameworks suggest a role for the ACC in surprise monitoring or optimizing cognitive control, respectively. So far, none of these frameworks has succeeded as a unified theory of ACC function. In this study, we reanalyzed previous neuroimaging data on ACC activity during cognitive tasks using a novel computational perspective: meta-Reinforcement Learning. We show that this computational framework can explain a variety of data on ACC function that, globally, could not be captured by any of the previous models. We propose that meta-Reinforcement Learning offers a unified theory of ACC cognitive and computational function.

## Introduction

Humans constantly face complex decisions, ranging from selecting one out of more available options (like choosing between an apple or a cupcake) to choosing the amount of cognitive and bodily resources we want to invest to achieve a goal (effort allocation). Formally, these decisional processes are aimed at solving a trade-off between minimizing the cost of investing cognitive (or physical) resources and maximizing the gain from investing a certain amount of resources^1–5^. In order to optimize this trade-off, it is sometimes necessary to expend some (cognitive) effort early in order to receive a larger reward later. In experimental psychology, the mental processes deputed to control the level of cognitive effort we deploy to achieve a goal are referred to as *cognitive control*. This process allows us to select more effortful options when we anticipate higher rewards from them. A typical example of cognitive control is the ability to inhibit habitual responses when these are not appropriate for the current situation (e.g. changing the route to your workplace when there are roadworks). The dorsal anterior cingulate cortex (dACC) and the surrounding cortical areas within the medial prefrontal cortex (MPFC) are known to be involved in cognitive control processes (e.g. ^6^), as well as in Reinforcement Learning (RL) and decision-making (e.g. ^7^), and the underlying computational mechanisms are still unclear, above all considering that their activation has been routinely found in many different experimental paradigms investigating several aspects of higher-order cognition^8–19^. Ironically, this area has been labelled as the Rorschach test for neuroscientists^20^.

The most recent and effective attempts to find a unified theory of dACC function have led to two competing frameworks: the Expected Value of Control framework^21^ (EVC) and the performance monitoring framework^8,22^. The EVC framework states that the dACC is involved in estimating the value of exerting *cognitive control* during a specific task and in selecting the optimal control signal, i.e. the control signal that maximises the estimated value^21^. This framework provides a theoretical explanation for several studies that show dACC activity during both anticipation of cognitively effortful tasks and the trade-off between the advantages of exerting cognitive control and its intrinsic cost ^18,23–25^.

The performance monitoring framework is based on RL and proposes that the dACC function is mainly aimed at performance monitoring by computing *prediction error* signals, resulting from the comparison between the expected and the actual outcome of actions. Prediction error signals are essential to update expectations about future action-outcomes associations. This theory was implemented in two independently developed models: the Predicted Response Outcome model (PRO) by Alexander & Brown^8^ and the Reward Value and Prediction Model (RVPM) by Silvetti et al.^22^ These models show that the combination of state-action-outcome expectations and the relative *surprise* signals (computed as the absolute value of prediction errors) was sufficient to explain many of the experimental findings about dACC function, from error detection to monitoring conflict between competing responses^26^.

Both the performance monitoring and EVC frameworks bear significant merits in offering mechanistic and integrative explanations of dACC functions, but none of them can explain the full range of empirical findings. A growing body of literature reports activity in the dACC linked to cognitive control (or more generally during anticipation of cognitive effort), in keeping with the perspective of the EVC framework^21^ (see also ^27,28^). However, given that in some of these cognitive control studies dACC activity is observed in the absence of surprise or prediction errors^3,24,29^ or while controlling for all possible sources of surprise^30,31^, these findings are not easily accommodated by the performance monitoring framework. Conversely, the performance monitoring framework provides a good account of the experimental findings documenting the role of the dACC in surprise coding during RL-based tasks^26^, while the EVC framework does not explain, by design, these results. Furthermore, a recent fMRI study compared the predictions of the two frameworks during a speeded decision-making task, which involved both cognitive control and performance monitoring^32^. This study suggests that the dACC function could be parsimoniously explained by performance monitoring mechanisms (involving prediction and prediction error), without postulating additional cognitive control optimization mechanisms, even in tasks requiring cognitive control^33^. Taken together, the conflicting results of the above studies create a theoretical impasse, since no single account seems to be able to provide an integrative account of dACC function that fully explains its multifarious roles in both cognitive control and performance monitoring.

Here, we show that an alternative theoretical framework overcomes this impasse by accounting for the roles of dACC in both cognitive control and performance monitoring: The Reinforcement Meta-Learner model^34,35^. The RML belongs to a framework named meta Reinforcement Learning (meta-RL), whose algorithms are able to adapt their internal parameters as a function of environmental challenges^36,37^. The RML combines some of the features of the EVC framework and the PRO model within the perspective of meta-RL, and in previous studies, it already provided a computational account of dACC function from both cognitive control and performance monitoring perspectives^34,35,38–40^.

In this study, we test the RML predictions about the dACC activity in two different experimental paradigms whose empirical results, taken as a whole, are irreconcilable with both the EVC and the monitoring frameworks. The first paradigm is the aforementioned speeded decision-making task, which provided support for the performance monitoring framework ^32^. The second paradigm is a verbal working memory task (WM) ^30,31,41^, showing dACC involvement in cognitive effort independently from surprise, in keeping with the perspective of the EVC framework.

We will show that the RML successfully simulates dACC activity both in the speeded decisionmaking task that provides support for models emphasizing performance monitoring and in the WM task that provides support for models emphasizing cognitive effort. These findings resolve the theoretical impasse generated by the competition between the two previous accounts of dACC, explaining its central role in both monitoring and cognitive control within an integrative perspective.

## Results

### RML description

The RML is an autonomous agent that learns and makes decisions (related to both motor behaviour and cognitive functions) to maximize net value, i.e. reward discounted by the cost of motor and cognitive effort (see Supplementary Information for a detailed description).

The model has been validated across multiple tasks and research domains ^34,35,38,39,42^, and was used in this work without parameter tuning (keeping all the parameters as in the original papers describing the model). From the neurophysiological perspective, the RML simulates a cortical-subcortical macrocircuit including the MPFC (including the dACC) and two brainstem neuromodulatory nuclei: the ventral tegmental area (VTA) and the locus coeruleus (LC) (Figure 1a). Previous literature has shown a connection between these areas (e.g. ^34,43,44^). Furthermore, the neurotransmitters produced by these brainstem nuclei, dopamine and noradrenaline, are known to affect decisionmaking (e.g. ^34,45,46^), and modulate the dACC function^47^.

**Figure 1.**
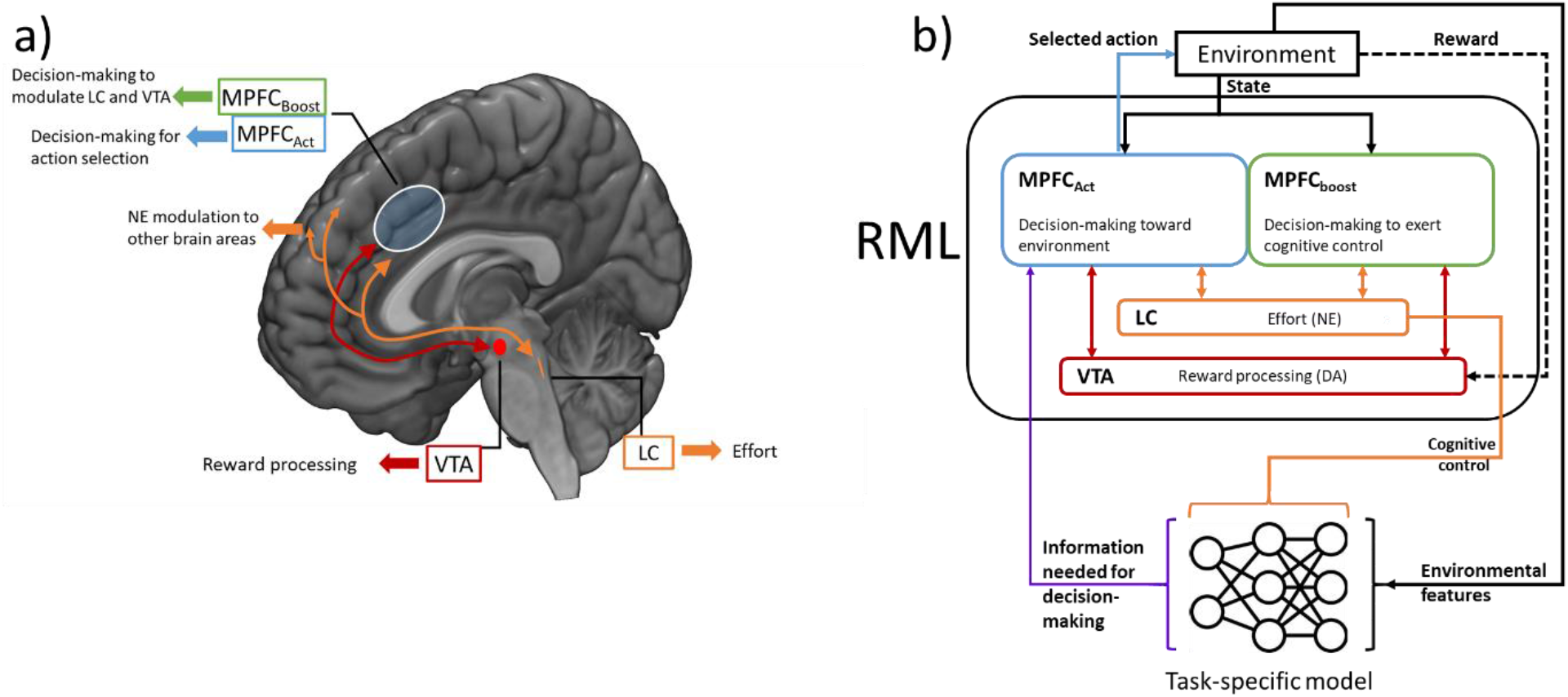
Description of the RML model. **a** Anatomo-functional mapping of RML modules. **b** Schematic representation of RML modules interactions. The agent consists of the RML and of a task-specific model that provides the RML with specific functions necessary to execute a task. The RML optimizes the task-specific model via LC output (that can be interpreted as the cognitive control signal).

The RML architecture assumes that the activity of the MPFC during decision-making can be explained by three different computations: the value of a decision (motor or cognitive), the surprise generated by the environmental feedback following this decision, and the level of cognitive control selected by the agent. Decision-making strategies relative to action selection and cognitive control levels (policies) are learned by agent-environment interaction, and based on approximate Bayesian learning implemented as Kalman filtering^48^. These three components of MPFC activity are processed by two separate modules (Figure 1b), an *action module (MPFC*_*act*_ *module)* – dedicated to motor action selection – and a *boost selection module (MPFC*_*boost*_ *module)* – dedicated to control optimization.

The MPFC_boost_ module performs control over motor and cognitive functions, by upregulating or downregulating activity in VTA and LC. When faced with a decision, the RML first decides on what amount of control to exert during the decision (termed ‘*boost’*). When the optimal boost level is selected, the MPFC_boost_ module sends a signal to the VTA and LC, which in turn influence both the MPFC modules. As the optimal policy for selecting the boost signal is learned, and the boost signal itself influences the learning process, we define the search for an optimal boost policy as meta-learning. Boosting the LC module promotes effortful motor actions and enhances information processing in other brain structures (cognitive effort) while boosting the VTA module improves reward signals and learning from non-primary rewards. Boost, however, implies an intrinsic cost, and the boost module implements meta-learning by dynamically optimizing the trade-off between performance improvement and boosting cost. In this work, we will equate the boost signal to cognitive control. This definition implies some simplification, as the boost signal can also influence the motivational component of decision-making, via VTA modulation, aside from cognitive and physical effort via LC modulation.

The MPFC_act_ module evaluates the options the agent has in the current environment based on expected effort and reward and selects the optimal action directed toward the environment. Action selection in the MPFC_act_ module is influenced by the LC input, which promotes effortful actions, and by the VTA input, which influences the expected value of actions. In this way, the action selection and value learning of the MPFC_act_ is indirectly modulated by the policy learned by the MPFC_*boost*_ module (meta-learning for optimal action selection).

The RML works as a task-independent optimizer of motor and cognitive decision-making. This is possible because the RML can be connected to external, task-specific, modules (e.g. a deep neural network; Figure 1b). The activity of the external module is controlled via LC signals, and its output is directed back toward the MPFC_act_ module to provide the RML with the perceptual and/or motor features needed for a specific task.

### Speeded decision-making task

#### Task description

We administered to the RML a simplified version of the speeded decision-making task proposed in Vassena et al. ^32^ (Figure 2a). In each trial, the agent was asked to select one among two options (representing fractal images in Figure 2a). The left and right options were independently set to yield a reward equal to an integer between 2 and 7 (six reward levels, with two fractal images for each of the six reward level). All possible combinations of the left and right reward were considered, for a total of 36 possible trial types. The goal of the agent was to select the best option (in terms of expected reward) in the shorter time possible. Once the RML selected the option, a reward corresponding to the fractal value was delivered to the model. In order to model the time pressure component of the task and incentivize the RML to respond as fast as possible, we implemented both a linear devaluation of the reward as a function of reaction time (supplementary Equation S13) and a response deadline. This solution captures the fact that human participants were instructed to respond as fast and as accurately as possible, and that no rewards were given in case of excessively long RT^32^. The difficulty of each trial was a function of the value difference between the presented options: large value differences led to faster and more accurate choices than small value differences.

**Figure 2.**
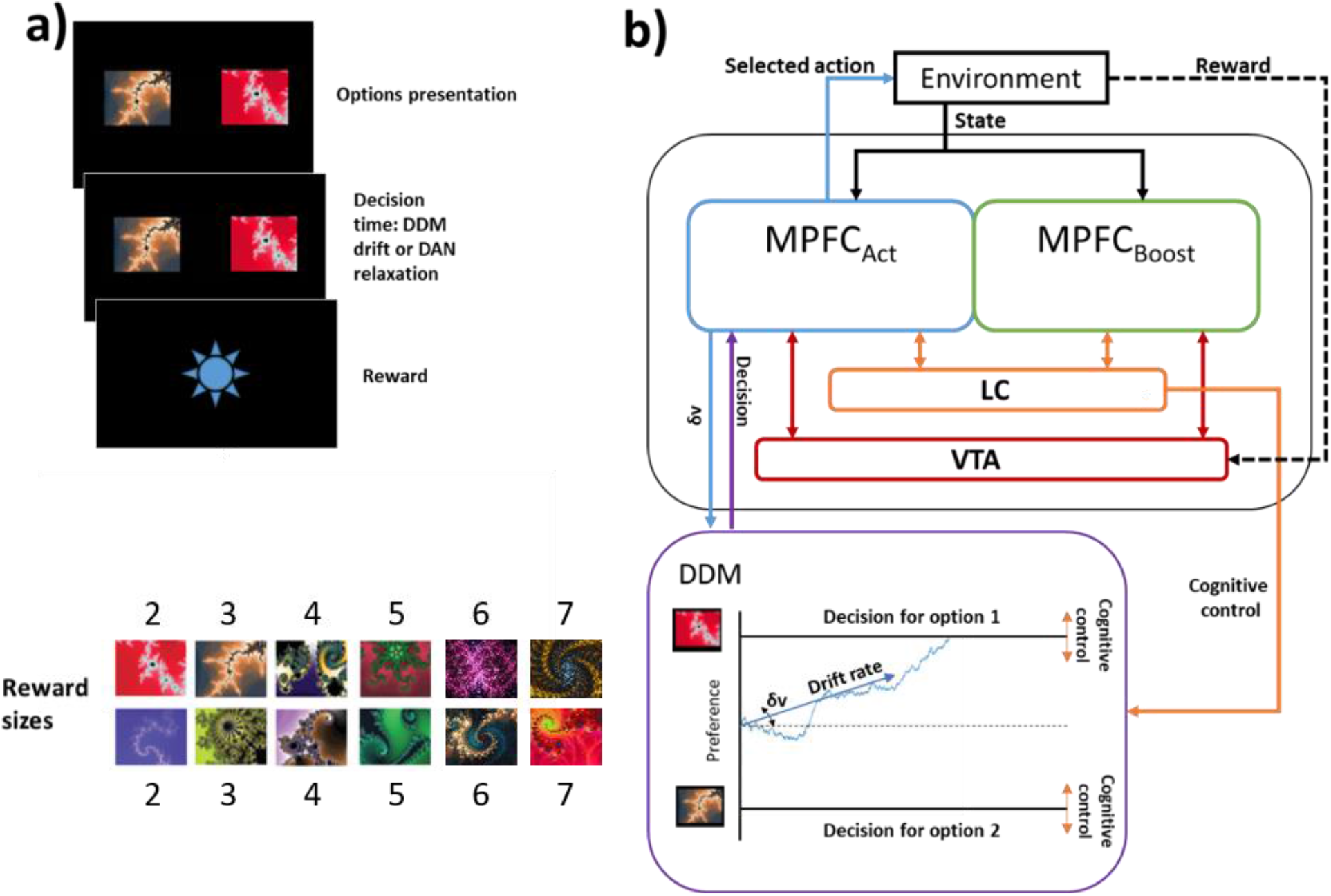
Speeded decision-making task: **a**: Setup of the speeded decision-making task, performed by the RML. **b**: The RML-DDM interaction during the speeded decision-making task. The RML receives input from the environment (about rewards and environmental states), and controls a task-specific module (the DDM), which helps in task execution. δv: difference in the expected value of the two options, whose absolute value determines the drift rate, while its sign determines the drift direction (up or down). The LC output modulated the decision boundaries.

#### The RML model in the speeded decision-making task

In order to make the RML able to perform the speeded decision-making task, we connected the model to an external module simulating decisions by accumulating evidence over time, therefore enabling the model to simulate reaction times. To show the generalizability of our results, we replicated our simulations with two different well-known models as external modules: the drift diffusion model (DDM) as proposed by Ratcliff^49^ and the dual attractor network (DAN) model by Usher & McClelland^50^. This ensured that the results could be attributed to the RML, and not to the specific external module used. In the main text, we describe the results of using the DDM as an external module. These were fully replicated with the DAN module, as described in detail in the supplementary material.

The integration of the DDM and the RML (Figure 2b) is inspired by Vassena et al.^32^, where the authors fitted the drift rate and the distance of decision boundaries of a DDM to their behavioural data to demonstrate that the speeded decision-making task relied on participants’ cognitive control. In each trial, the DDM received two different signals from the RML as an input. The first was the difference in the expected value of the options (*δv* in Figure 2b) from the MPFC_act_ module. The value of each possible option was learned by the RML during a training session preceding the task. The absolute value of the *δv* signal determined the DDM drift rate, while the sign of the *δv* signal determined the drift direction toward one of the decision boundaries (representing the two available options, left vs right option) for that trial. In this way, a larger *δv* absolute value resulted in higher accuracy and faster decision^32^. The second input from the RML to the DDM consisted of the LC signal, which modulated the distance of the decision boundaries. The higher the LC activity, the closer the boundaries, and the faster the decision. As the LC output is controlled by the boost signal from the MPFC_boost_ (cognitive control), the latter can therefore modulate the decision speed as a function of trial type. Once the DDM reached one decision boundary, or no decision was made within the response time, this information was passed to the MPFC_act_ module that implemented the option selection; finally, both the MPFC modules updated the expected value for the selected option based on the environmental feedback (reward).

#### Simulation results

In Figure 3a we report the dACC activity measured with fMRI during the speeded decision-making task, as reported in Vassena et al.^32^. This can be described as a W-shaped function of the difference in value between the two options (quartic function with a positive leading coefficient). During the execution of the same task, the RML predicted the W-shaped activity pattern exhibited by the dACC (Figure 3b). Using the Akaike information criterion (AIC)^51^, we found that the simulated dACC activity follows a quartic pattern with a positive leading coefficient rather than a quadratic pattern (Aikake weight for a quartic function > 0.999), similar to Vassena et al.^32^

**Figure 3.**
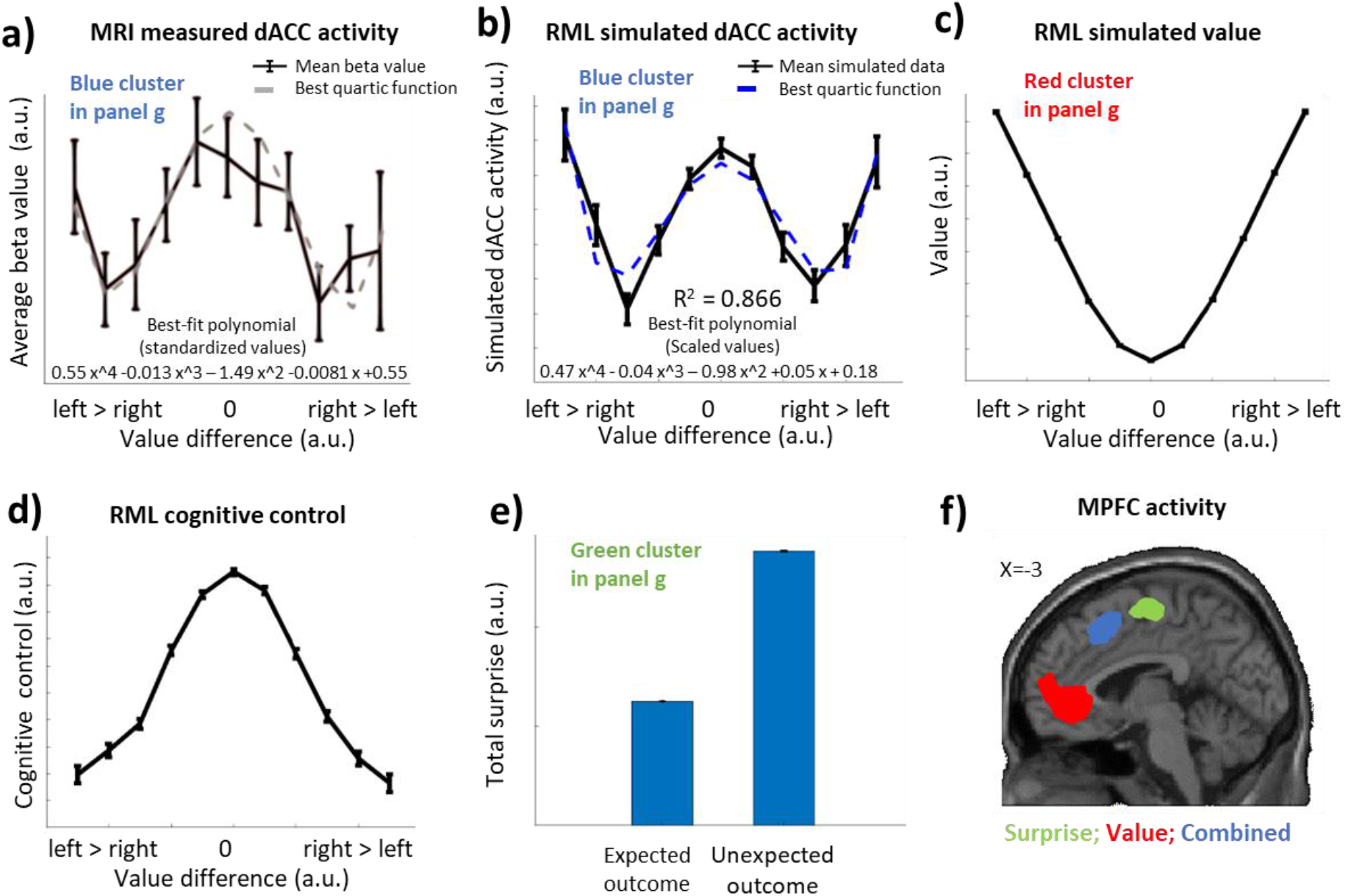
Results of the speeded decision-making task. **a:** The MRI results from Vassena et al.^32^ (adapted). The dACC activity is shown in the black line, while the grey, dashed line shows the best fitting quartic function to this data. **b**: The dACC activity as simulated by the RML (black line), and the best fitting quartic function (blue, dashed line). This activity is the combination of the value (panel c) and the cognitive control (panel d). **c**: The value component of the RML activity. **d**: Cognitive control signal (RML boost) as a function of stimuli value difference. **e:** RML surprise-related activity. **f:** Activation clusters within the MPFC (adapted from Vassena et al.^32^). Blue: dACC activation as a mixture of value and cognitive control, RML prediction in panel b; Red: vMPFC value-based activation, RML prediction in panel c; Green: mid-cingulate activation relative to average surprise, RML prediction in panel e.

The RML simulates the W-shaped dACC activity - shown in Figure 3b - as the combination of two different neural signals evoked by the cue onset: the expected net value (expected reward discounted by expected costs, Figure 3c) and the cognitive control signal (boost signal in the RML, Figure 3d).

The expected net value component is dependent on the difference in cue value and follows a U-shaped function. When the values of both options are similar (around 0 on the x-axis in Figure 3c), the mean expected value of that trial type is minimal, while it is maximal for large value differences. Two mechanisms cause the shape of this function. First, similar option values lead to smaller drift rates in the DDM, and therefore to longer RTs (see also Equations S10, S13). Since long RTs cause reward devaluation and a higher probability to exceed the response deadline, the RML learns that the execution of this type of trial has a lower average value (minimum in the U-shaped pattern in Figure 3c). Conversely, when the difference between option values is large, RTs are shorter on average (higher drift rates), leading to a higher expected value (maxima in Figure 3c). The second mechanism depends on the intrinsic cost of boosting. The boost function mirrors the net value function (inverted U shape, see below), and the boost cost function – which is a linear function of boost - follows the same shape. For this reason, the expected value is maximally discounted in the centre and minimally discounted in the tails, contributing to a U-shaped expected value. Figure 3d shows the contribution to the RML simulation of dACC activity given by the cognitive control signal (boost), which follows an inverted U-shaped function. As the boosting level maximizes reward while minimizing the cost intrinsic to boosting, the RML increases the boost only if the consequent reward gain is worth the boost cost. When the difference between the options is small, the DDM drift rate is small. In that case, a larger boost allows for a significant improvement in performance by shrinking the decision boundaries and therefore shortening the RTs (Figure 2b, see also equation S11 in the supplementary materials). Conversely, when the task is easy, and the value difference is large, the DDM drift rate is high. In that case, RTs will be fast anyway, and increasing the boost will only marginally improve them. This leads to a reduced boost signal (cognitive control) when the value difference is large (minima in Figure 3d).

The RML explains the W-shaped dACC activity equally well as the PRO model, which was successfully used in the original study ^32^. Moreover, only the RML provides a further experimental prediction about another region of the MPFC: The vMPFC. Indeed, the value function in Figure 3d predicted the activity of this region (red cluster in Figure 3f), which is well known to be associated with value estimation^52,53^. Finally, the RML also predicts the dACC activity evoked by the reward-driven surprise computed as the absolute difference between the overall reward available in a trial and the long-term average reward (Figure 3e and green cluster in Figure 3f). This result is due to the surprise-coding mechanisms proper of both the RML and the PRO models.

### Working memory task

#### Task description

In this simulation, we administered a WM task like the one used in Engström et al. ^31^ (Figure 4a), to the RML. This task exemplifies the role of the dACC in cognitive effort. During each trial, the RML was exposed to either 1, 4, 6 or 8 words, generating four different difficulty levels. After a delay of 10s, the model was presented with a target word that matched one of the memorized words in 50% of trials. The model’s goal was to indicate whether the target word matched one of the words presented before.

**Figure 4.**
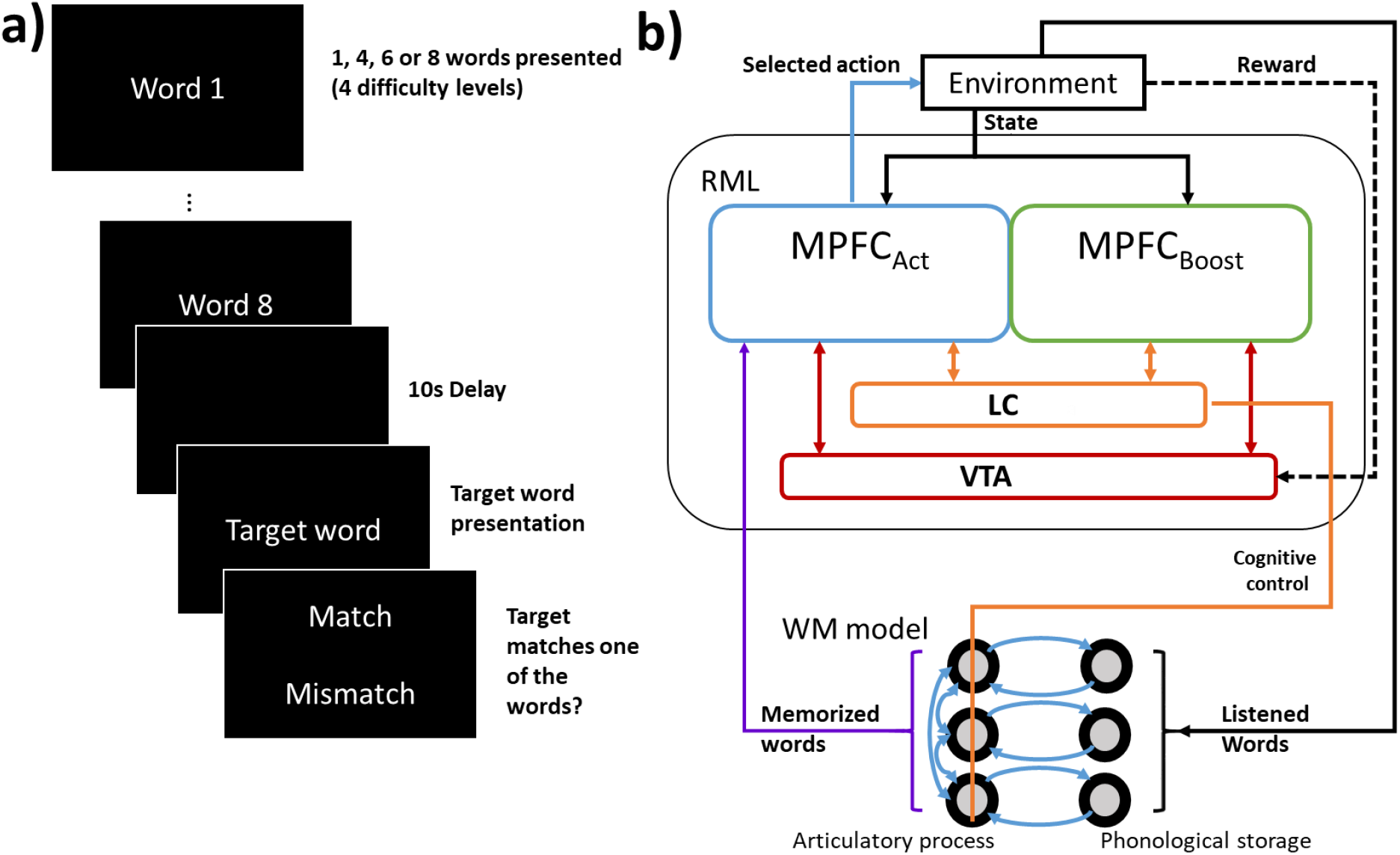
Verbal WM task. **a**. Setup of the verbal WM task, performed by the RML. **b**. Schematic representation of RML-cRNN interaction during the execution of the verbal WM task. The LC output modulated the gain of the neural units in the articulatory process layer, improving words retention in WM.

#### The RML model in the WM task

For this task, the RML was connected to a task-specific model that simulated verbal WM functions (Figure 4b). This model consisted of a dual layer competitive recurrent neural network (cRNN), inspired by the FROST network^54^. The input layer of the cRNN encoded the words presented in each trial (each unit encoded one word), working as a phonological storage. The output layer retained the words during the delay period, thanks to recurrent connectivity with the input layer. This layer simulated the articulatory process and also compared the memory content with the target word. Precision of words retention decreased with increasing WM load (number of words to be remembered) due to lateral inhibitory connections in the output layer. The RML improved words retention by gain modulation of the output layer, via LC output. The MPFC_Act_ made the decision about words-target matching based on the linear combination of the cRNN output (see Supplementary Methods for details). Here we ran 15 simulations, representing 15 different subjects.

#### Simulation results

In Figure 5a we show the dACC activity measured with fMRI during the WM task. As reported in Engstrom et al.^31^, the dACC activity was significantly described by a linear function of WM load. The RML successfully predicted the dACC activity (Figure 5b) as boost intensity by the MPFC_Boost_ module, showing –like in the fMRI reults-a significant linear trend as a function of WM load (t-test on beta values: t(14) = 17.58, p < 0.0001). This result derives from the optimization of the boost signal as a function of task difficulty, in order to counteract –via LC output-the cRNN drop of performance due to increasing WM load.

**Figure 5.**
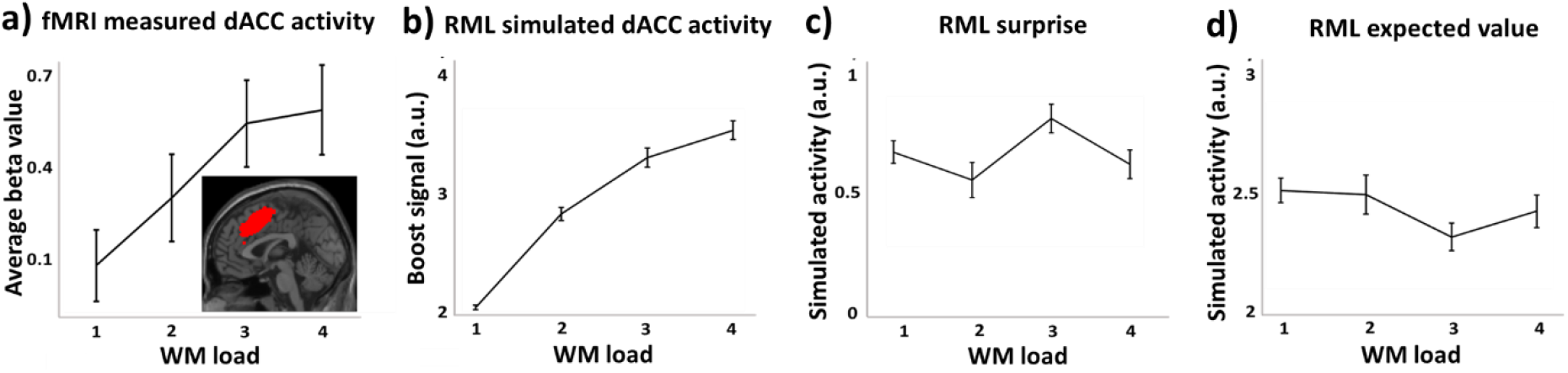
Results from verbal WM task. **a)** fMRI results from Engstrom et al. ^31^ showing dACC activity (red cluster in subpanel) as a function of WM load (difficulty levels 1-4). **b)** Boost signal from RML as a function of WM load. **c)** RML surprise (unsigned PE, average of MPFC_Act_ and MPFC_Boost_) as a function of WM load. **d)** RML expected value (average of MPFC_Act_ and MPFC_Boost_) as a function of WM load.

The RML also showed that neither surprise (Figure 5c) nor value expectation (from both MPFC_Act_ and MPFC_Boost_, Figure 5d) can explain the dACC activity during the task (none resulted to show a significant linear trend as a function of WM load). This was because accuracy remained constant across difficulty levels (mean accuracy 87%, main effect of WM load: F(3,14)= 1.39, p = n.s.), like reported in human participants ^41^.

## Discussion

In this study, we proposed the meta-RL framework as a solution to the theoretical impasse generated by fact that the two most prominent accounts of dACC in human cognition – namely, the performance monitoring framework and the EVC framework – focus on two distinct sets of studies, whose results are difficult to reconcile within a unified account. While the performance monitoring framework (instantiated with the PRO model) was effective in predicting the dACC activity in a cognitive control task involving unexpected events—the speeded decision-making task utilized in Vassena et al.^32^,—it is ineffective in predicting the dACC activity in cognitive control tasks designed to exclude unexpected events (e.g. ^3,24,29–31^). Differently, the EVC perspective could successfully frame the aspects of the dACC activity related to cognitive control optimization in the absence of surprise signals but does not account for the neural data when surprise and unexpected events are relevant for task execution. ^7^

Here we have shown that the meta-RL perspective (instantiated with the RML model^34,35^) can explain dACC activity in both task types, as it grounds cognitive control optimization on Bayesian surprise tracking. Our two simulations showed that the meta-RL perspective can simultaneously account for findings that were previously explained separately by the EVC and PRO models. When applied to a speeded decision-making task that exemplifies the importance of surprise monitoring, the RML matched the PRO model in predicting the dACC activation (Figure 3b) and surprise-driven activity in the mid-cingulate cortex (Figure 3e), while also additionally predicting the vMPFC activity related to value estimation (Figure 3c), found by Vassena et al.^32^ as a replication of experimental results from previous literature (e.g. ^52,53^). When applied to a working memory task that exemplifies the importance of cognitive effort, the RML successfully predicted dACC activity (Figure 5b) and the fact that neither surprise (Figure 5c) nor value expectation can explain the dACC activity during the task (Figure 5d).

It is important to note that while both the RML and PRO models generate comparable predictions regarding dACC activity in the speeded decision-making task ^32^, they provide distinct underlying computational explanations. The PRO model generates the W-shaped dACC activity pattern as the combination of two different types of surprise signals. One surprise signal is triggered by the cue onset, and it depends on the uncertainty relative to the cue type. The other is triggered by the response onset and depends on the uncertainty related to the probability of selecting one of the two responses over the other. The PRO model therefore suggests that the dACC plays no role in the optimization of cognitive control per se but it is exclusively involved in surprise tracking. As argued above, this view is challenged by empirical evidence showing dACC activation corresponding to cognitive control intensity in tasks deliberately designed to exclude surprise^3,24,29–31^. In contrast, the RML explains the dACC activity as the sum of two different functions: The expected value of the selected action (U-shaped function in Figure 3c) and the cognitive control intensity (inverted U-shaped function in Figure 3d), suggesting a more complex function of the dACC, where surprise-based monitoring is functional to cognitive control optimization.

This is the reason why the RML, differently from the PRO model, is able to explain the dACC activity also in tasks where surprise is not involved, such as the working memory task studied here (see also ^34,35,38^). These findings solidify the RML as a computational account unifying the monitoring and control perspective during decision-making, thereby reconciling surprise, value, and cognitive control under the framework of meta-learning processes^36,37^.

Another important innovation of the RML is its bio-inspired system-level perspective, where MPFC function is studied in conjunction with brainstem and midbrain catecholaminergic nuclei. This allows for modeling of the cortico-subcortical reciprocal influence and how the MPFC can orchestrate cortical and striatal functions by controlling the release of neuromodulators, broadening the scope of application of the RML framework. For example, in a previous work, the RML was shown to be capable of providing a computational account of how the MPFC modulates parietal neurons - via LC activation - during a visual attention task^35^. The same mechanism can also explain the activity in dorsolateral regions of the frontal and parietal lobes that Vassena et al. described in their study^32^. Indeed, these areas are involved in the execution of the speeded decision-making task (spatial and motor control components), and the RML suggests that the MPFC optimizes their performance by neuronal gain modulation via neuromodulatory input. Moreover, this system-level perspective allows us to formulate predictions about different regions within the MPFC, like the activity pattern of the vMPFC (Figure 3c) or of the mid-cingulate cortex (Figure 3f).

A further relevant element introduced by the RML is that most of the behavioural and neural predictions made by the RML emerge from the interaction of the model with the environment (*situatedness* as defined by Wilson^55^ and Nolfi^56^). Furthermore, the goal of the optimization process of the RML is to find the set of actions (policy) that maximizes the net value, regardless of whether these actions are directed toward the external environment (motor) or the internal environment (cognitive control). Learning optimal actions (motor or cognitive) requires a loop between the RML and the environment, where the RML plays an *active* role in exploring the world. Recently, an improvement to the RML was proposed, the RML-C^35^, where the situated and proactive aspects of RML are more pronounced, as the model’s goal is not limited to net value maximization, but includes also the maximization of information relative to the environment. To this aim, the RML-C actively explores the environment to improve its predictions, investing motor and cognitive effort for gathering information (*intrinsic motivation*^57,58^), beyond the utilitarian perspective of net value maximization. The situated and proactive perspective that the RML proposes about the dACC role – and more in general of the MPFC – seems to be soundly grounded also on the neurophysiology of this area, which evidences large neural populations coding for both motor and visceromotor functions (for a review: ^26^). Differently from the RML, the PRO model embraces a pure monitoring perspective, where the dACC plays a passive role of RL “critic” ^59^, dedicated to the monitoring of action-outcome contingencies. Similarly, the implementation of the EVC model proposed by Vassena et al.^32^ selects the control intensity as a function of the option values and the control cost, working as a decision-making algorithm where information about the environment and the action-outcome contingencies is fully known in advance. However, a more recent implementation of the EVC, the Learned Value of Control (LVOC) model^28^, has been designed for learning the optimal policies of control through model-environment interactions in stationary conditions. However, contrary to the RML and PRO models, the LVOC does not include explicit monitoring of surprise, hence it remains to be tested whether it could account for the dACC data relative to surprise coding.

In summary, in this work, we have shown that the meta-learning perspective represents a general solution for the understanding of the MPFC function and that it is capable of explaining empirical data from a larger domain set if compared with the PRO and EVC models alone. Future research in this domain should investigate also the comparison of the above frameworks with other, alternative perspectives on the computational basis of cognitive control, based on active inference^60,61^, such as the proposal that cognitive control results from the monitoring of the deviation from prior beliefs about cognitive actions so that cognitive effort is exerted to override habits^62^.

## Methods

A detailed description of simulation methods and data analysis is reported in the Supplementary Information.

## Acknowledgments

TV is a PhD student enrolled in the National PhD in Artificial Intelligence, XXXVII cycle, course on Health and life sciences, hosted by Università Campus Bio-Medico di Roma. TV is funded by national grant PRIN 2022, grant N° 20227MPSEH. MS is funded by national grant PRIN 2022, grant N° 20227MPSEH.

