## Supplementary information for "Meta-Reinforcement Learning reconciles surprise, value, and control in the anterior cingulate cortex"

### Supplementary methods

#### A detailed description of the RML

In this study, we model decision-making using a computational model, the reinforcement meta-learner (RML[1]). While already summarized in the main text, this section aims to expand on the description of the RML, and use formal descriptions of the processes described in the text. As the aim is to provide an explanation that can be read without prior knowledge on this model, several of the formulas provided in this document come from an earlier study describing the RML[2]. For any further clarifications, we will refer to this paper.

As a reinforcement learning algorithm, the RML models decision-making as a Markov decision process in discrete time. It assumes that the environment over time can be split in a sequence of states, denoted by $s$. The state at a certain timepoint $t$ is denoted by $s_t$. In each of these states, the agent is prompted to select one of several different actions. Each of these actions, denoted by $a$, all have action-outcome contingencies. Each action will result in both a state transition to state $s'$, and a direct reward, denoted by $r$. We note that this direct reward can be zero, meaning that an action only changes the current state of the environment, without yielding a reward. Importantly, the model does not know these contingencies at the start of the task: they have to be learned.

The RML models the action selection and learning as two distinct parts of an action. Once a state is observed, an action has to be taken, and only after this action is taken, and the outcome is received, can the value of this action be updated.

The RML has several features distinguishing it from other reinforcement learning paradigms. First, it takes two dimensions of actions into account. Their expected value, and the amount of effort – be it physical or mental – required to execute this action. In order to select the most optimal value, the reward received from the action is discounted by the effort required to obtain it. The optimal action is then selected using a softmax function.

A second feature is its ability to regulate several of its own internal parameters. For one, the RML regulates the extent to which the reward is discounted by the effort. Additionally, the reward sizes are regulated.

Thirdly, the RML incorporates meta-learning; it is able to learn about the learning itself. The RML is able to optimize its own learning rate based on the volatility of the environment, enabling the agent to optimize the speed at which it learns.

The RML achieves this using three different modules. It uses a **boost module**, an **action selection module**, and a **control module**. Both the **boost module** and **action selection module** perform action selection in a way similar to an actor module in traditional reinforcement learning paradigms. The **boost module** regulates the optimization of the effort discounting and reward size parameter. It does so by selecting a boost intensity, indicating to what extend the model should upregulate its internal parameters. Critically, this upregulation of parameters is costly. The higher the effort discounting and reward size parameter, the higher the cost of the boosting action. The action selection module selects the most optimal action to be executed by the agent. Finally, the **control module** has two functions. It communicates the signal from the **boost module** to the **action selection module**, and it evaluates the outcome of the selected action, allowing the **boost module** and **action selection module** to learn about the outcome of their selected actions.

In the next part, we will first explain the equations used by the RML to optimize decisions. Afterwards, we give a detailed description on the neural structure underlying the decision-making process by the RML.

#### Action selection

The RML models action selection in two steps. First, the optimal intensity of boosting is selected. This boosting has two functions: it regulates the effort discounting and reward size parameter, as detailed below. This is performed by the **boost module**. The **boost module** selects the optimal boosting using the perceived environmental state. In the RML, this intensity of boosting is discrete: several predetermined intensities are available to be selected. This allows for the boost module to work in a way similar to a Q-learning actor, where the possible actions are the intensities of boosting available (the intensity of boosting at timepoint $t$ is denoted by $b_t$). When faced with an environmental state, the boost module selects a value for $b_t$ by evaluating the expected reward for each option from a stored table denoted by $V_{boost}(b_t)$. Then, the realized boost intensity is selected using a softmax function (see Equation S1).

This intensity of boosting does not directly influence the **action selection module**. First, the **boost module** sends the signal for the selected intensity of boosting to the **control module**. The **control module** then transforms this *boost* value using a linear equation (see Equation S2). This transformed value is sent to the **action selection module**, where it is used to make a decision about which action to select. To this end, the **action selection module** considers both the current state $s_t$, the possible actions ($a_1, \ldots, a_n$), and the learned action values (denoted by $V_{act}(a_t)$). It selects an action using a softmax, as described in Equation S3.

**Learning**

After this action is selected and executed, the agent receives feedback from the environment. This feedback can come in two different forms: a reward (denoted by $R$), and a transition to a different state (denoted by $s'$). When this feedback is received, the **control module** processes this feedback into an experienced value, specific to the **action selection module** (denoted by $DA_{act}$, see Equation S4a). Additionally, it calculates the experienced value for the **boost module** (denoted by $DA_{boost}$, see Equation S4b).

In the **action selection module**, the experienced value is the sum of the received value and the expected future reward, where the boost value selected during the action selection increases this value linearly. Furthermore, this value takes both the immediate expected value of the action, and the expected value of future actions into account. In the case of the **boost module**, the experienced reward is the actual realized reward in the trial, including the expected value of future rewards in this state, discounted by a temporal discounting factor, where the cost of boosting is subtracted. Since the RML hypothesizes that a higher boost level is more costly, this cost of boosting is linearly dependent on the boost intensity $b$.

The model uses these experienced values to calculate the prediction error (PE), the difference between the expected value and the experienced value. This PE is then used to update the expected value for the selected action $a$ (see Equation S5a) and boost intensity $b$ (see Equation S5b).

**Dynamic learning rate**

Contrary to other reinforcement learning paradigms, the RML hypothesizes that the learning rate for the **boost module** and **action module** are dynamic, and dependent on the environment. The learning rates increase when the RML encounters a large change in environment, to quickly learn the new values encountered in the environment. When the agent encounters small PE values, the learning rates decrease, to minimize the effect a single large PE has on the learned values. The RML hypothesizes that this calculation can be modeled using a Kalman filter[3].

For the **action selection module**, the learning is based on the action PE, i.e. the difference between the experienced action value and the expected action value based on earlier experience (see equations S5a). The RML calculates the PE, and models the learning rate as an approximate Kalman filter to estimate the volatility of the environment. To this end, the expected average variance of the value is divided by the squared estimated PE (Equation S6). The learning rate is set to a minimum value of $\beta$, to ensure some learning from each trial, avoiding a situation where the agent gets stuck in a state where they cannot learn anymore.

Both the estimated variance and the estimated prediction error are modeled as timecourses, and smoothed over time. For the estimated variance, a smoothed value timecourse is generated first, which is defined recursively based on the value of the smoothed value in the previous timecourse, as a linear combination of the value in the previous timepoint and the current value (equations S7 and

S8). Similarly, the prediction error is smoothed over time, where the absolute value of the current prediction error is used instead of the signed prediction error (Equation S9).

In a similar way, the RML estimates the dynamic learning rate for the **boost module**, using the boost PE instead of the action PE (see Equation S5b). The calculation of the Kalman filter is performed similarly to the action learning rate, the difference being that the RML keeps track of the variance, time smoothed value, and time smoothed PE for the **boost module** separately from the **action selection module**.

**RML-to-brain mapping**

The RML is a bio-inspired system based on the meta-reinforcement learning framework proposed by Doya et al.[4] It is inspired by previously identified bidirectional connections between different neural structures, specifically the medial prefrontal cortex (MPFC), locus coeruleus (LC), and ventral tegmental area (VTA) (e.g. [5–7]). Both the **action selection module** and **boost module** are hypothesized to simulate MPFC, including the dorsal anterior cingulate cortex (dACC)[1,2]. These modules are bidirectionally connected with the **control module**, which is represented by several brainstem nuclei (specifically the LC and VTA). These brainstem nuclei perform the different tasks that are prescribed to the **control module** in the above text. The VTA generates a dopamine (DA) signal, which provides an experienced reward signal to the **action selection module** and the **boost module**. The effort discounting parameter is signaled by the LC, by a generated noradrenaline (NE) signal. Additionally, the before mentioned adaptive learning rate is tracked in the LC. Finally, one of the strengths of the RML is its flexibility: we are easily able to connect the RML structure described here to an **external module**. This external module can be influenced by the activity in the **control module** and the expected value of the different actions, and in turn influence the decisions made by the RML by affecting the **action module**. See Figure 1a and 1b in the main text for a graphical representation of the anatomical representation and connections of these modules.

#### RML Equations

$$p(selected\ boost = \ b|s) = \sigma(v_{\text{boost}}(s,b), \tau_{boost}) \tag{S1}$$

This equation defines the probability *p* of selecting boost level *b* while in state *s*. $v_{boost}(s,b)$ is the expected value of boost level *b* given the current state *s*. $\tau_{boost}$ indicates the temperature of the softmax function $\sigma$.

$$NE_{LC}(b) = f(b) \tag{S2}$$

In this equation, $NE_{LC}(b)$ is the LC signal (i.e. the LC-generated *NE* signal), which is set as a function dependent on the boost level *b*. In the paper by Silvetti et al.[1] describing the RML, this formula is set to be the identity, for the sake of simplicity. Therefore, the value of $NE_{LC}(b)$ is set to be equal to the boost value.

$$p(chosen\ action = a|s) = \sigma(v_{act,t}(s,a) - \frac{C(s,a)}{NE_{LC}(b)}, \tau_{act}) \tag{S3}$$

This equation determines the probability *p* of selecting a certain action *a,* directed toward the environment, given state *s*. $v_{act,t}(s,a)$ is the learned value of action *a* given state *s*. $\tau_{act}$ indicates the temperature of the softmax function $\sigma$. $C(s,a)$ is the cost value of action *a* given state *s*.

$$DA_{act,t} = (r_t(R_t + \mu b) + b(1-\mu)\rho \max_{a \epsilon A_{s'}}(v_{act,t}(s',a))) \tag{S4a}$$

$$DA_{boost,t} = r_t R_t - \omega b + \max_{b\epsilon B_{s'}}(v_{boost,t}(s', b)))$$ (S4b)

Equations S4a and S4b show the calculation of the VTA signals (i.e. the VTA-generated DA signals) for both the action (S4a) and boost (S4b) module. For the action module, this signal is a combination of the presence ($r_t$) and size ($R_t$) of the reward at time *t*, with an added value depending on the amount of cognitive control, in case a reward was present. This cognitive control is multiplied by a meta-parameter of the system governing the DA dynamics, $\mu$. Furthermore, the most optimal future reward (i.e. the maximum reward of any action in the next state) is taken into account, multiplied with a temporal discounting parameter $\rho$. In the equation, $A_{s'}$ is the collection of all possible actions given state *s'*.

For the boost module, the VTA signal is defined as the actual reward received in the trial, where the cost of the boosting is subtracted. The cost of the boosting is equal to the level of boosting, multiplied with $\omega$, a parameter describing the cost of boosting. The higher this parameter, the more costly it is to perform boosting. Additionally, the boost module also looks at the most optimal future reward by estimating the optimal reward in the next state.

$$\Delta v_{act,t}(s, a) = \lambda_{act,t} \cdot (DA_{act,t} - v_{act,t-1}(s, a))$$ (S5a)

$$\Delta v_{boost,t}(s, b) = \lambda_{boost,t} \cdot (DA_{boost,t} - v_{boost,t-1}(s, b))$$ (S5b)

Equations S5a and S5b show the update of the learned expected state-action value after a trial. The learning system uses the dynamically changing learning rate ($\lambda_{act,t}$ and $\lambda_{boost,t}$ respectively), as calculated in Equation S6, as well as the experienced value of the trial at time *t*, conveyed by the VTA-generated DA signal ($DA_{act,t}$ and $DA_{boost,t}$ respectively) to update its internal value. After calculating the update value ($\Delta v_{act,t}(s, a)$ or $\Delta v_{boost,t}(s, b)$), the expected value is updated by adding the update value.

$$\lambda_{act,t} = \max\left(\frac{\widehat{Var_t}(v)}{\hat{\delta}_t^{\,2}}, \beta\right)$$ (S6)

This equation calculates the learning rate for the action module as the quotient between the expected variance of *v* at timestep *t* ($\widehat{Var_t}(v)$, calculated in Equation S7), and the squared prediction error at timestep *t* ($\hat{\delta}_t^{\,2}$, calculated in Equation S9). If this value is smaller than $\beta$, the learning rate is set to $\beta$ to avoid computational instability. As described in previous papers[1,2], the equations S6-S9 approximate a Kalman gain[3].

$$\widehat{Var_t}(v) = (v_t - \hat{v}_{t-1})^2$$ (S7)

Equation S7 describes the generation of the expected variance of *v* at time *t* ($\widehat{Var_t}(v)$), as a function of the timecourse of the value $v_t$ and the time-smoothed value at the previous timestep ($\hat{v}_{t-1}$), calculated in Equation S8.

$$\hat{v}_t = \hat{v}_{t-1} + \alpha \cdot (v_t - \hat{v}_{t-1})$$ (S8)

Equation S8 describes the timecourse of the time-smoothed value $\hat{v}_t$: each timestep, it updates as a combination of the old value of the time-smoothed value, and the difference between the current value of the trial and the old time-smoothed value, with a weight of $\alpha$. This $\alpha$ is a parameter determining the amount of smoothing in the time-smoothed value, as well as the amount of smoothing in the time-smoothed prediction error (as shown in Equation S9).

$$\hat{\delta}_t = \hat{\delta}_{t-1} + \alpha \cdot (|\delta_t| - \hat{\delta}_{t-1}) \quad \text{(S9)}$$

Equation S9 shows the calculation of the time-smoothed prediction error ($\hat{\delta}_t$), which is defined similarly to the time-smoothed value in Equation S8. It is a combination of the old value of the time-smoothed prediction error, combined with the difference between the absolute value of the current prediction error and the old value of the time-smoothed prediction error, with a weight $\alpha$.

| Free parameters | | | |
|---|---|---|---|
| Variable | Description | Equation | Value in simulation |
| $\tau_{boost}$ | Temperature of the softmax for the boost module | S1 | 0.3 |
| $\tau_{act}*$ | Temperature of the softmax for the action selection module | S3 | N.A. |
| $\mu$ | Parameter indicating DA dynamics | S4a | 0.1 |
| $\rho$ | Temporal discounting parameter | S4a | 0.1 |
| $\alpha$ | Temporal smoothing parameter in determining learning rates | S8, S9 | 0.3 |
| $\beta$ | Minimum learning rate | S6 | 0.15 |

*Supplementary Table 1: variables used in the RML, and how they are found. Although these parameters are indicated as free parameters in the original RML, they are fixed in the current study; we did not vary these parameters during this study to optimize the fit between the actual behavior and the simulated behavior. The final column indicates the values of the parameters as used in the simulations reported in the main text and supplementary methods, inherited from [1].*

#### Speeded decision-making task modeling

In order to model the task by Vassena et al.[8] using the RML, we defined 36 possible states. Each of these states consists of a two-option decision-making task. The left and right options are independently set to yield a reward equal to an integer between 2 and 7. All possible combinations of the left and right reward are considered. Each state has an intrinsic difficulty, defined by the difference between the reward received from the left and right reward. We modeled the task difficulty inversely linear to the difference between the expected value of the left and right option; the more similar both options are, the more difficult it is to make a decision between them.

Before modeling the task, the RML is given a prior value for each set of fractals, equal to the actual value of the fractal. Then, the RML is given an explorative training, where it explores all options and decisions, in order to optimize the prior value for each set of fractals. Afterwards, the RML performs 1944 trials (54 trials per state). During the analysis, these states are sorted by value difference (the difference between the left and right option), and the z-scored mean boost and sum of the mean value of the boost and actions are plotted. The total dACC activity is set to the sum of these two values. In total, 200 simulations are performed, and the average results are reported.

For this task, we implemented two separate external modules, the drift diffusion model, and the dual attractor model (described in detail below).

##### The drift diffusion model

The drift diffusion model (DDM) was first introduced by Ratcliff[9]. It models a decision between two options by keeping track of a confidence variable, indicating both which option the agent is currently favoring, and to what extend the agent is confident about its choice. The model accumulates its confidence over time, until a preset decision boundary is reached. At that point, it selects the option corresponding to the decision boundary reached.

Mathematically, the confidence generated during each timestep is a combination of the drift rate (indicating the average confidence gain due to the environment, denoted by *DR* in this paper to avoid confusion with the variable indicating value in the RML) and random Gaussian noise (with a mean of 0, and standard deviation of $\sigma$). Normally, the DDM uses two decision boundaries. One occurs when the accumulated confidence value reaches a threshold level (denoted by $\theta'$), and the other when it reaches 0. In an unbiased version of the DDM, the accumulated confidence starts at a value of $0.5 \cdot \theta'$ at the beginning of each trial. For simplicity, we make a small notational change to this model. Instead of starting at $0.5 \cdot \theta'$, and performing a decision at either 0 or $\theta'$, we let the model start at 0, and make a decision when the confidence reaches either - $\theta$ or $\theta$, where $\theta$ is equivalent to $0.5 \cdot \theta'$.

In our combination, we combine the RML with the DDM to gain two different outputs. First, we are interested in which decision the DDM makes, indicated by the decision boundary reached by the confidence variable. Additionally, we are interested in the duration it took to reach a decision, which can be used to determine the reaction time of the trial.

**RML-DDM interface**

In order to combine the DDM with the RML, several RML values are used as input in the DDM (see Figure 2 from the main text for a visual overview of the connection between the DDM and the RML). Specifically, the value of the drift rate and threshold level both depend on RML parameters. The drift rate is directly proportional to the expected difficulty of the trial, the difference in the expected value of one option compared to the other. It is set according to Equation S10:

$$DR = v_{DDM} \cdot (v_{act}(a_1, s) - v_{act}(a_2, s)) \tag{S10}$$

Here, $DR$ is the drift rate, $v_{DDM}$ is a model parameter that determines the influence of the difficulty on the drift rate. $v_{act}(a_1, s)$ and $v_{act}(a_2, s)$ denote the estimated value selecting action 1 or action 2 in the current state *s* as computed by the RML respectively.

Additionally, the initial threshold is set to the quotient of the boost level and a model parameter, scaling this boost level for use in the DDM, as shown in Equation S11:

$$\theta = \theta_{DDM}/b \tag{S11}$$

Here, $\theta_{DDM}$ is a model parameter, while *b* is the boost level during the current trial.

The DDM starts by setting the confidence value (denoted by $c(t_{TR})$, a function of $t_{TR}$, the time passed within the trial) to 0, at time $t_{TR}$=0 and evolves over time following Equation S12, until $c(t_{TR})$ is either equal to $\theta$, or $-\theta$. Additionally, in case the model has not reached either decision threshold when the time limit is reached, the model is set to be too late in making a decision. In this case, no reward is given in this trial.

$$c(t_{TR} + dt_{TR}) = c(t_{TR}) + \left(DR + N(0, \sigma_{DDM})\right) \cdot dt_{TR} \tag{S12}$$

This equation describes the evolution of *c* over time. In this equation, *DR* denotes the drift rate for the current trial (calculated according to Equation S10), and $N(0, \sigma_{DDM})$ denotes random noise drawn from a normal distribution with mean 0 and standard deviation $\sigma_{DDM}$, a model parameter of the DDM.

The DDM then provides the RML with both the decision made and the value of $t_{TR}$ at which this decision is made. This value of $t_{TR}$ is used to get the reaction time. In order to ensure the RML optimizes its decisions to optimize both accuracy and reaction time, the model receives a reward penalty depending on its reaction time (see Equation S13). The slower the agent responds, the larger this penalty is.

$$R = RW_{act} - \frac{RT_{DDM}}{v_{RT}} \tag{S13}$$

In this equation, the trial reward ($R$) is a function of the received reward due to the chosen action ($RW_{act}$), and the penalty based on the reaction time. This penalty is given as the quotient between the found reaction time of this trial due to making a decision according to the DDM ($RT_{DDM}$) and a model parameter ($v_{RT}$) indicating to what extent the DDM-based *RT* influences the final received reward.

##### Free parameters of the DDM

This implementation of the combination of DDM and RML has several model parameters that can be seen as free parameters. These parameters are changed in our simulations to determine what model parameter values lead to the optimal estimation of the data by Vassena et al.[8] In this study, we did not change the parameters in the RML from Silvetti et al.[1], but only varied several of the indicated parameters that govern the combination of the DDM and the RML. These parameters are summarized in supplementary table 2. In total, four free parameters are estimated. The first parameter indicates to what extent the difference in expected value as estimated by the RML influences the DDM drift rate. A second parameter is used to indicate to what extent the boost is scaled to get the DDM threshold. A third parameter is used to determine the standard error of the Gaussian noise in the DDM. The final parameter is used to discount the RT output from the DDM, and convert this to a reward penalty.

| DDM parameters | | | |
|---|---|---|---|
| Parameter | Parameter description | Equation | Value used |
| $v_{DDM}$ | Scaling factor for the value difference in the DDM | S10 | 1.598 |
| $\theta_{DDM}$ | Scaling factor for the threshold in the DDM | S11 | 14.41 |
| $\sigma_{DDM}$ | Standard error of the Gaussian noise in the DDM | S12 | 71.99 |
| $v_{RT}$ | Scaling factor for the reaction time output | S13 | 62.04 |

*Supplementary Table 2: the free parameters varied in the combined RML and DDM model, and the optimal values.*

##### Optimization of free parameters for the DDM

In order to find the optimal values of the free parameters, we performed a gradient descent (GD) procedure on all four free parameters. The loss function of this GD procedure was a combination of the mean squared error of the accuracy and the response time outputs by the RML-DDM combined model compared to the real accuracy and response times reported by Vassena et al.[8] Here, accuracy is defined as the percentage of simulated trials the agent made the optimal decision (i.e. the decision leading to the highest reward). As the response time output by the DDM is unitless, and would only indicate the variable part of the response time (the non-decision time is not taken into account in our simulations), the RT was first z-scored before being added to the loss function. The parameter values corresponding to the optimal GD output are reported in the final column of supplementary table 2.

##### The Dual attractor network

In order to show the results attained using the RML-DDM combination are not solely due to the influence of the DDM, we implemented a second model. For this second model, we combined the RML with the dual attractor network (DAN), described by Usher & McClelland[10]. Contrarily to the DDM, the DAN models decision-making not as a competition of one information variable, where two different boundaries determine which action is chosen, but using two different variables. These variables represent the information accumulation for both actions (i.e. the certainty the action is optimal). While these different variables (attractors) increase independently based on the information available in the environment, they are connected through mutual competition: each of these attractor states decreases the value of the other state proportional to its value. A decision is made when one of the two of these parameter values reaches a pre-set threshold.

This decision-making is modeled by accumulating information about both options over time. Each timestep, the information grows depending on the input, the perceived difference in information.

Additionally, the attractors are considered to be leaky; with each timestep, the information decreases by a proportion of the current information. Finally, the attractors are competing. The information for one attractor state directly decreases the information for the other attractor state (see equations S14a and S14b, adapted from Usher & McClelland[10], with some changes in notation to avoid confusion with parameters from the RML).

$$x_1(t_{TR} + dt_{TR}) = x_1(t_{TR}) + dt_{TR} \cdot (I(1) - \kappa \cdot x_1(t_{TR}) - \beta_{DAN} \cdot x_2(t_{TR}) + \xi) \qquad \text{(S14a)}$$
$$x_2(t_{TR} + dt_{TR}) = x_2(t_{TR}) + dt_{TR} \cdot (I(2) - \kappa \cdot x_2(t_{TR}) - \beta_{DAN} \cdot x_1(t_{TR}) + \xi) \qquad \text{(S14b)}$$

In this equation, $x(t_{TR})$ indicates the information for both options at the time, where $x_1(t_{TR})$ indicates the accumulated information for option 1, while $x_1(t_{TR})$ indicates the accumulated information for option 2. This information is updated using the information from the previous timestep, adding the input for options 1 and 2 ($I(1)$ and $I(2)$ respectively). From this value, part of the current value is subtracted (the decay of information, with a characteristic decay rate parameter $\kappa$). In this study, $\kappa$ is a model parameter, which is estimated during the parameter optimization step detailed below. Furthermore, the collected information for the other option influences the information for the current option; this information is multiplied with $\beta_{DAN}$ (renamed from the original equation by Usher & McClelland to avoid confusion with the $\beta$ parameter defined in the RML), a parameter indicating the strength of the inhibitory connection between the options, and subtracted from the information. Finally, random Gaussian noise ($\xi$) is added, with standard deviation $\sigma_{DAN}$). In the simulations, $dt_{TR}$ is set to 1/10.

**RML-DAN interface**

In our current study, we combine the DAN with the RML. In this combination, several parameters from the RML are used as input for the DAN. These are used by the DAN to generate both a selected action and a reaction time of this choice, which are provided to the RML. Similar to the DDM implementation, the amount of cognitive control and the expected value of both actions are given as parameters to the DAN (see Supplementary Figure 1 for a visual representation of the combination between the DAN and the RML). The amount of cognitive control influences the strength of the inhibitory connection between the options ($\beta_{DAN}$ in equations S14a and S14b), while the difference in expected value influences the difference in information gain between the actions ($I$ in equations S14a and S14b).

The transformation of the level of cognitive control to the inhibitory connection strength is given in Equation S15: $\beta_{DAN}$ is set to the amount of cognitive control $b$, divided by 10, multiplied with a model parameter ($Dual_{inf}$). Additionally, the difference in expected value between both options is used to determine the information gain parameter. The information gain parameter difference is set to the difference in value between options 1 and 2, multiplied with a model parameter ($Dual_{cong}$) (Equation S16a and S16b). As in Usher & McClelland[10], this value is added to 0.5 and subtracted from 0.5 in order to ensure a total information gain of 1.

$$\beta_{DAN} = b/10 \cdot Dual_{inf} \qquad \text{(S15)}$$

In this equation, b represents the level of boosting, which is brought to a value between 0 and 1 by dividing it by 10.

$$I(1) = 0.5 + Dual_{cong} \cdot (v_{act}(1) - v_{act}(2)) \qquad \text{(S16a)}$$

$$I(2) = 0.5 - Dual_{cong} \cdot (v_{act}(1) - v_{act}(2)) \qquad \text{(S16b)}$$

In these equations, $v_{act}(1)$ and $v_{act}(2)$ represent the expected values of selecting options 1 and 2 according to the RML respectively.

The DAN then determines both the selected action and the value of $t_{TR}$ at the moment the action is selected and provides these values back to the RML. The value of $t_{TR}$ is then converted to the response time, and in the same way as with the RML-DDM combination, this response time is used to add a penalty to the received reward (see Equation S17). The slower the agent responds, the larger this penalty is.

$$R = RW_{act} - \frac{RT_{DAN}}{v_{RTDAN}} \quad \text{(S17)}$$

In this equation, the trial reward (R) is a function of the received reward due to the chosen action ($RW_{act}$), and the penalty based on the response time. This penalty is found by dividing the found reaction time of this trial due to making a decision according to the DAN ($RT_{DAN}$) by a model parameter ($v_{RTDAN}$) indicating to what extent the DAN-based RT influences the final received reward.

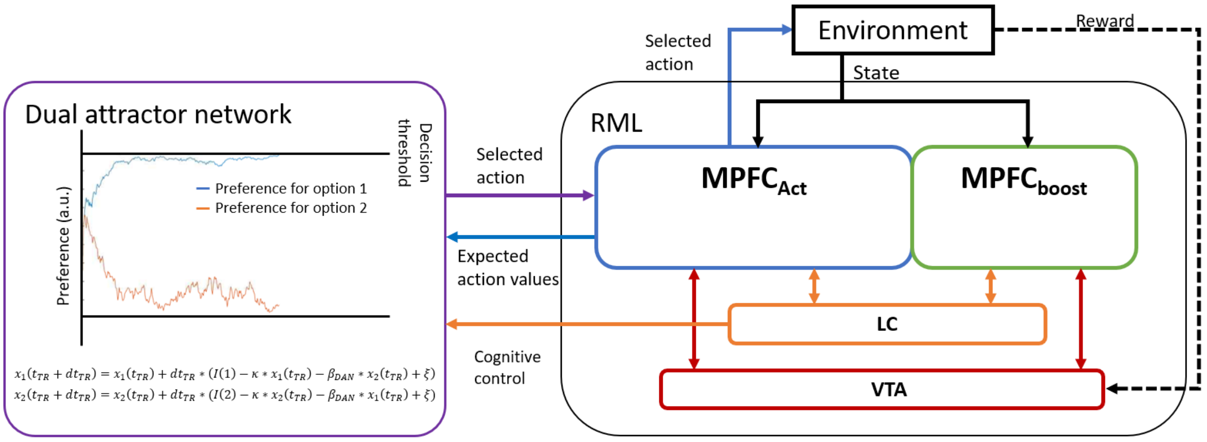


***Supplementary Figure 1****: Visual representation of the used connection between the RML and the DAN. As with the DDM representation, the DAN has a bidirectional connection to the action module of the dACC. On the one hand, the expected action values are taken from the dACC, and sent to the DAN. The difference between these action values is set as the difficulty of the trial. This value influences the external information gain (I) in each step in the dual attractor framework. This information gain is advantaged for the option with the higher expected reward. On the other hand, the DAN generates an action and response time, which are sent back to the action module of the dACC. There, they are combined with the received reward, and the RML updates the internal values for this action. Additionally, the DAN receives input from the LC. The amount of cognitive control determines the strength of the competition between both attractor states. A higher boost value would therefore allow an increase in information for one condition to inhibit the other condition stronger.*

#### Free parameters of the DAN

This implementation of the combination of DAN and RML has several model parameters that can be seen as free parameters. These parameters are changed in our simulations to determine what model parameter values lead to the optimal estimation of the data by Vassena et al.[8] As mentioned before, we did not change the parameters in the RML from Silvetti et al.[1], but only varied several of the indicated parameters that govern the combination of the DAN and the RML. These parameters are summarized in supplementary table 3. In total, five different free parameters are estimated. These are two free parameters in the update equations of the DAN (equations S14a and S14b), one for the characteristic decay rate, and one for the standard deviation of the Gaussian noise. Additionally, one scaling parameter is used for the value difference input, and one for the boost input. A fifth free parameter is used to convert the RT output from the DAN to a reward. These parameters are optimized (as described below), and the optimal values are shown in the final column of supplementary table 3.

| DAN parameters | | | |
|---|---|---|---|
| Parameter | Parameter description | Equation | Value used |

| | | | |
|---|---|---|---|
| $\kappa$ | Characteristic decay rate | S14a, S14b | 0.408 |
| $\sigma_{DA}$ | Standard deviation of the Gaussian noise $\xi$ | S14a, S14b | 0.284 |
| $Dual_{inf}$ | Scaling factor for the information in the DAN | S15 | 0.727 |
| $Dual_{cong}$ | Scaling factor for the congruency in the DAN | S16a, S16b | 0.017 |
| $v_{RTDAN}$ | Scaling factor for the reaction time output | S17 | 99.63 |

*Supplementary Table 3: the free parameters varied in the combined RML and DAN model, and the found optimal values.*

**Optimization of the DAN free parameters**

Similar to the DDM, the DAN free parameters are estimated using a GD procedure varying all five free parameters. The loss function of this GD procedure was a combination of the mean squared error of the accuracy and the response time outputs by the RML-DAN combined model compared to the real accuracy and response times reported by Vassena et al.[8] As before, accuracy is defined as the percentage of simulated trials the agent made the optimal decision (i.e. the decision leading to the highest reward). As the response time output by the DAN is unitless, and would only indicate the variable part of the response time (the non-decision time is not taken into account in our simulations), the *RT* was first z-scored before being added to the loss function. The parameter values corresponding to the optimal GD output are reported in the final column of supplementary table 3.

#### Verbal working memory task modeling

During each trial (Figure 4a, main text), 1, 4, 6 or 8 words were presented to the model, generating four different difficulty levels. After a delay of 10s, the model was presented with a target word that matched one of the memorized words in 50% of trials. The model's goal was to indicate whether the target word matched one of the words presented before. In case of correct response, the model received a reward signal equal to 3, while it received no reward for an incorrect response. The RML first performed a training session, consisting of 40 trials for each difficulty level (160 trials). Afterwards, it performed the task, consisting of 90 trials for each difficulty level (360 trials in total), randomly intermixed. We repeated the simulation 20 times (simulating 20 participants). RML parameters were the same as in the original paper [1] (Supplementary Table 1).

**Working memory (WM) model**

The RML was connected to a task-specific external module (Figure 4b, main text) simulating items encoding and maintenance in the WM [11]. This module consisted of a two-layered competitive recurrent neural network (cRNN). The input layer (16 units) encoded the words (both target and words to be memorized), while the output layer (16 units) maintained the words, thanks to recurrent connectivity with the input layer. We assigned arbitrarily one input unit to each word. Each output unit was connected with symmetrical weights to one input unit. Output units were globally connected with symmetric lateral inhibitory weights. We arbitrarily assigned a duration of 10 ms for each network update cycle. For this reason, for example, a delay of 10s meant 1000 network cycles. The lateral inhibitory connections in the output layer ensured an overall decrease of activity as a function of the number of active units, simulating the detrimental effect of increasing WM load on item retention [11]. Equations and a more detailed description of the cRNN, including the parameters set we used in this study, can be found in the Supplementary Material of our previous study [1].

**RML-cRNN interface**

The RML-cRNN interface from our previous study was designed for a dynamical implementation of the RML[1]. Here, we redesigned the interface, as in this study we used an MDP implementation of the RML. The activity of the cRNN output layer was modulated by the NE signal from the LC module of the RML (Equation S2), defining a variable *h* indicating a preference for selecting the 'match' action, based on the activity of each neuron in the output layer of the cRNN (Equation S18):

$$h = NE_{LC} \cdot \max(F) - \epsilon \quad \text{(S18)}$$

Where $F$ is the vector of the cRNN output layer activity, $h$ is a scalar resulting from the activity modulated by the LC output $NE_{LC}$ subtracted by a threshold parameter $\epsilon = 0.15$. When $h > 0$, the RML has more evidence for a match trial, while a value of $h < 0$ indicates that the RML has more evidence for a mismatch trial. For each trial, action selection about the presence of a match is performed by Equation S3, selecting between the 'match' action (indicating a match is present), and the 'mismatch' action (indicating a match is absent), via modulation of the state-action values as stated in equations S19a and S19b.:

$$v^*_{act}(s, a_{match}) = v_{act}(s, a_{match}) + \varphi \cdot h \quad \text{(S19a)}$$

$$v^*_{act}(s, a_{mismatch}) = v_{act}(s, a_{mismatch}) - \varphi \cdot h \quad \text{(S19b)}$$

Where $v^*_{act}$ is the cRNN-modulated state-action value (to be used in Equation S3), $a_{match}$ indicates the index of the 'match' choice, $a_{mismatch}$ indicates the index of the 'mismatch' choice, $\varphi = 50$ is a scaling parameter, and $h$ is the preference parameter set in Equation S18.

#### Supplementary results

In this section, we describe the results of the RML-DAN combined model analysis (see Supplementary Figure 2). Furthermore, although most of our free parameters are involved in the way the RML and the DDM or DAN communicate, one of the free parameters controls the degree to which a slower response contributes to the reward (the $v_{RT}$ and $v_{RTDAN}$ parameters for the DDM and DAN respectively). To investigate the effect a change in this parameter has on the simulated dACC activity, and therefore measure the robustness of the simulations, we redid the simulations for different values of this parameter (see Supplementary Figure 3).

**Speeded decision-making task with RML-DAN combined model**

The results from the RML-DAN combined model are shown in Supplementary Figure 2. In Supplementary Figure 2a, we have shown the dACC activity as measured by Vassena et al.[8], in order to compare this activity to the simulated dACC activity by the RML-DAN combined model (shown in Supplementary Figure 2b). Similar to the results of the RML-DDM combined model, we can see that the simulated dACC activity follows a W-shaped pattern (the dashed blue line in Supplementary Figure 2b), closely reproducing the found dACC pattern. After using the AIC to test the most likely order of the simulated dACC activity (as described by Wagenmakers & Farell[12]), we find that the simulated dACC activity is more likely to be a quartic function of the value difference rather than a quadratic one with a positive leading coefficient (Akaike weight > 0.999, equivalent to a p-value < 0.001). This dACC activity is the sum of the mean expected value (Supplementary Figure 2c), and mean cognitive control (Supplementary Figure 2d). The mean expected value shows (Supplementary Figure 2c), like in the RML-DDM combined model, a u-shaped function. We can see an increase in expected value with a larger value difference. The minimum in the expected value when the value difference is close to 0 can be explained in two ways. First, when the value difference is small, the difference in information gain is small (see Equation S16a and 16b). This leads to a situation where the accumulated information for both options is equal. Since the accumulated information for one option inhibits the information for the other option (see Equations S14a and S14b), this causes the *RT* to increase, in turn causing a larger penalty term (see Equation S17), and thus a lower value. Secondly, a higher value of boosting causes a larger cost due to the intrinsic cost of boosting. Since a lower value difference increases the boost value (as detailed below), these trials yield a lower value compared to the trials where there is a large difference in value. The mean cognitive control function (Supplementary Figure 2d) shows a similar shape to the mean cognitive control function in the RML-DDM combined model: an inverted U-shaped function. Since boost carries an intrinsic cost, the boost is only increased when necessary, yielding an increase in boost when the trial is difficult (i.e. when the difference between the options is small). In these cases, an increase in boost level lowers the inhibition between the two accumulators (see Equation S15), allowing for a faster *RT*. This has two advantages: first, it directly reduces the penalty term to the reward. Second, it allows the RML-DAN combined model to respond in time in relatively more trials.

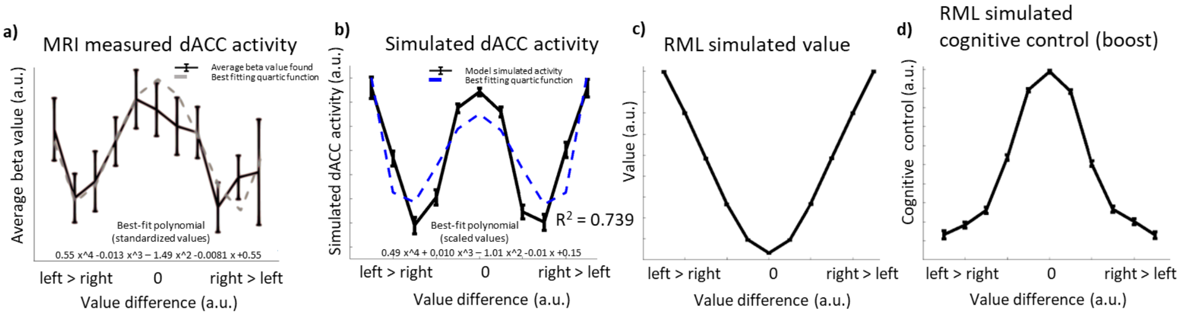


***Supplementary Figure 2**: Results from the DAN combined with the RML.* ***a**: The real dACC activity measured using fMRI by Vassena et al.*[8] ***b:** Solid line: the dACC activity as simulated using the RML combined with the DAN. This activity is a combination of the expected value and cognitive control simulated by the RML. The dashed line is the best-fitting quadratic function. We can see that similar to the fMRI data, the quartic function fitting the simulated dACC activity has a positive first coefficient.* ***c:** The expected value simulated for different value differences. This expected value shows a u-shaped function, with a maximum for the extreme value differences.* ***d:** The simulated cognitive control level for different value differences. Contrary to the value function, the cognitive control shows an inverted u-shaped function, with a maximum value when the value difference is 0.*

**Effect of changes in the $v_{RT}$ and $v_{RTDAN}$ parameter on the simulated dACC activity**

Besides a change in the parameters governing the interface between the RML and the DDM or DAN, one of the parameters we base our model inversion on is the parameter governing how the amount of steps the model takes is converted into a reward devaluation for the model. In order to investigate the stability of the dACC simulations with changes in this parameter, we have performed the simulation for several different values of either $v_{RT}$ (in case of the DDM) or $v_{RTDAN}$ (in case of the DAN), changing the value from the optimum by a certain percentage between an addition of 75%, and a subtraction of 75%. Supplementary Figure 3 shows the effects of a change in this parameter. The RML-DDM combined model shows mild effects on the simulated dACC activity after a change in the $v_{RT}$ parameter. The RML-DAN combined model does show some differences in dACC activity when changing the $v_{RTDAN}$ parameter. Although an increase does not affect the shape of the simulated dACC activity function too much, a large decrease (75%) shows a change from a quartic function to a sixth-degree polynomial function.

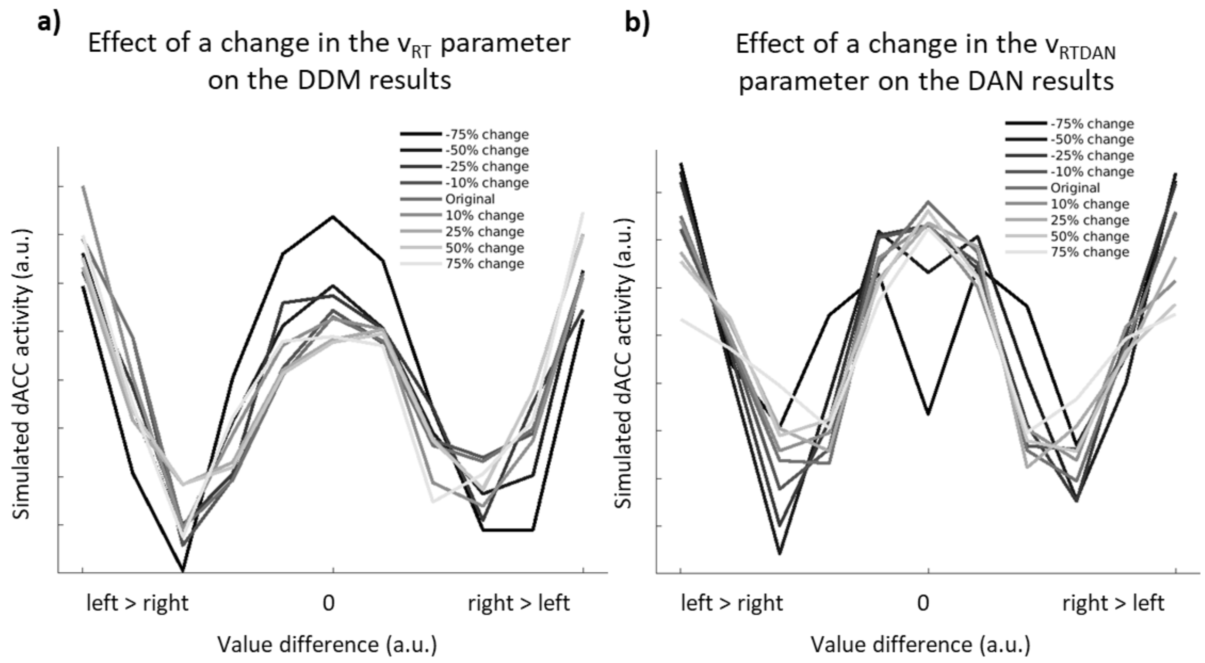


***Supplementary Figure 3**: **a)** Simulated dACC activity is mildly influenced by different levels of time pressure to respond ($v_{RT}$ parameter in Equation S13). Original: $v_{RT}$ value used in the simulation shown in the main text (Suppl. Table 2). Variations as a percentage of the original value. Results from the RML-DDM simulations. **b)** Same as in a), but relative to the RML-DAN simulations. Original: $v_{RTDAN}$ value reported in Suppl. Table 3 and used in Equation S17.*
